# Computing the Orientational-Average of Diffusion-Weighted MRI Signals: A Comparison of Different Techniques

**DOI:** 10.1101/2020.11.18.388272

**Authors:** Maryam Afzali, Hans Knutsson, Evren Özarslan, Derek K Jones

## Abstract

Numerous applications in diffusion MRI, from multi-compartment modeling to power-law analyses, involves computing the orientationally-averaged diffusion-weighted signal. Most approaches implicitly assume, for a given b-value, that the gradient sampling vectors are uniformly distributed on a sphere (or ‘shell’), computing the orientationally-averaged signal through simple arithmetic averaging. One challenge with this approach is that not all acquisition schemes have gradient sampling vectors distributed over perfect spheres (either by design, or due to gradient non-linearities). To ameliorate this challenge, alternative averaging methods include: weighted signal averaging; spherical harmonic representation of the signal in each shell; and using Mean Apparent Propagator MRI (MAP-MRI) to derive a three-dimensional signal representation and estimate its ‘isotropic part’. This latter approach can be applied to all q-space sampling schemes, making it suitable for multi-shell acquisitions when unwanted gradient non-linearities are present.

Here, these different methods are compared under different signal-to-noise (SNR) realizations. With sufficiently dense sampling points (61points per shell), and isotropically-distributed sampling vectors, all methods give comparable results, (accuracy of MAP-MRI-based estimates being slightly higher albeit with slightly elevated bias as b-value increases). As the SNR and number of data points per shell are reduced, MAP-MRI-based approaches give pronounced improvements in accuracy over the other methods.

## Introduction

Diffusion MRI is a non-invasive technique that is sensitive to differences in tissue microstructure, which comprises a combination of micro-environments with potentially different orientational characteristics. Inherent to MRI is averaging of the magnetization across the voxel. However, some orientational features (macroscopic or ensemble anisotropy) typically survive such averaging, making applications like fiber-orientation mapping feasible. As far as studies aiming to understand the underlying microstructure are concerned, the presence of such macroscopic anisotropy may introduce complications in the interpretation of the signal. In analogy with solid-state NMR applications^1^, considering the “powdered” structure of the specimen that features replicas of each and every microscopic domain oriented along all possible directions could more clearly reveal the desired microstructural properties of the medium^2,3^.

To factor out the effect of macroscopic anisotropy in diffusion MRI, i.e., to estimate the signal for the “powdered” structure, two approaches have been proposed: (i) taking the “isotropic component” of the signal^3,4^; this is typically achieved by representing the signal with a series of spherical harmonics and keeping the leading term; and (ii) numerical computation of the orientational average of the diffusion-weighted signal profile^5,6^. Due to the emerging interest^7–16^ in employing the so-called “powder-averaged signal” for tissue characterization, we consider the problem of estimating this quantity from data acquired via common single diffusion encoding (SDE) protocols.

The accuracy of the powder-averaged signal depends on both the set of gradient directions employed in the data acquisition and the numerical method used to estimate the average. Regarding the former, different strategies have been proposed to optimize the sampling strategy, the most well-known and widely-used of which is the electrostatic repulsion algorithm^17^. In this approach, a uniform single-shell distribution of *q*-space sample points is found by minimizing the electrostatic energy of a system containing antipodal charge pairs on the surface of a sphere. Situations under which such single-shell diffusion sampling schemes are rotationally-invariant have been discussed previously^18,19^. In some applications, the sample points should be optimally distributed not just on a single sphere, but across three-dimensional *q*-space^20^. Knutsson et al.^21^ proposed a novel framework that extends the traditional electrostatic repulsion approach to generate optimized three-dimensional *q*-space sample distributions, by enabling a user-specific definition of the three-dimensional space to be sampled and associated distance metric.

The second important factor in orientational signal averaging is the method used to compute the powder-average, which is our focus in this study. Methods that perform brute-force estimation of the signal average as well as those that take the “isotropic component” of the signal, (derived from signal representations), are compared.

## Results

### Effect of noise and number of samples

Figure 1 illustrates the results obtained from 61 × 8 samples for both shelled and non-shelled point sets in the presence of Gaussian noise. We note that in this sampling scheme^21^, we do not have the orientationally-averaged signal from the Lebedev method because this weighted-averaging approach requires its own point sets and weights. Figure 1(a) shows the mean and standard deviation of the estimated signal versus b-value using the MAP-MRI method^22^ with *N*_max_ = 6 for five different noise standard deviations, *σ*_*g*_ = 0.1414, 0.0707, 0.0283, 0.0071, 0.0014, and three different dispersions, *κ* = 1, 9, ∞. The mean and standard deviation of the *d*_1_ and *d*_2_ measures are illustrated as, respectively, a dot and an error bar in Figure 1(b) for each estimation method.

**Figure 1.**
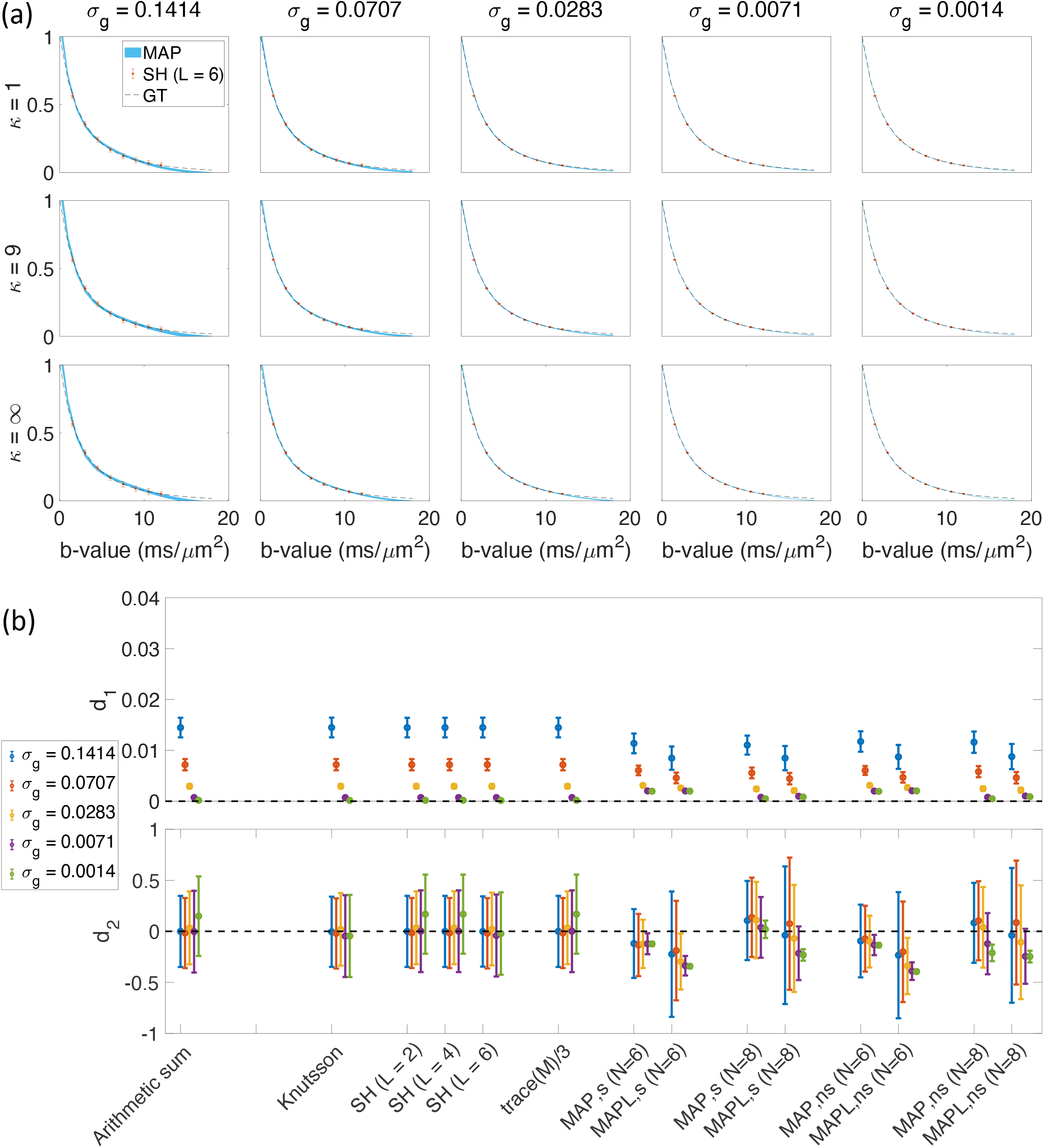
The results from 488 samples for both shelled (61 × 8) and non-shelled point sets^21^ in the presence of Gaussian noise. (a) the mean and std of the estimated signal versus b-value using MAP-MRI method with *N*_max_ = 6 for five different noise floors, *σ*_*g*_, and three different dispersion values, *κ*. The thickness of the blue band is twice the standard deviation of the signal estimates and its center is the mean. The dashed black line shows the ground truth and the red dots and bars show the results of the SH (L = 6), spherical harmonic representation. (b) the mean and std of the *d*_1_ and *d*_2_ measures for different methods. **Arithmetic sum:** simple arithmetic averaging; **Lebedev:** weighted averaging by^26^; **Knutsson:** weighted averaging by^27^; **SH:** Spherical harmonic method for powder averaging by^3^ *L* = 2, 4, 6, shows the order in spherical harmonic representation; **trace(M)/3:** powder average signal from equation (10); **MAP:** direction-averaged signal using MAP-MRI^22^ for *N*_max_ = 6 and 8; and **MAPL:** direction-averaged signal using MAP-MRI with Laplacian regularization^48^. The ‘s’ and ‘ns’ correspond to the shelled and non-shelled point sets, respectively. Different colors (blue, red, yellow, purple, green) show the results in different noise levels (sg = 0:1414; 0:0707; 0:0283; 0:0071; 0:0014).

The error, quantified by *d*_1_, is small for all methods. Under noisy conditions, MAP-based methods outperform the shell-by-shell estimates, and the regularization employed in MAPL method evidently yields a further reduction in *d*_1_. At very low noise levels, for *σ*_*g*_ ≤ 0.0071, *d*_1_ is very small for all methods, though slightly elevated for MAP and MAPL methods with *N*_max_ = 6. The *b*-dependent bias is quantified through *d*_2_. The mean *d*_2_ values are generally better in shell-by-shell estimates. However, the standard deviation of *d*_2_ is large in almost all the shell-based methods even for very low noise levels (*σ*_*g*_ = 0.0014), which is likely a side-effect of treating shells independently in the averaging schemes. However, the standard deviation of *d*_2_ is substantially reduced in MAP-based methods as the noise level decreases. As far as bias is concerned, MAP performs better than MAPL in most cases.

Figure 2 shows the results of 344 (43 × 8) samples in the presence of Gaussian noise while Figure 3 illustrates the same for the sampling schemes with 152 (19 × 8) samples in the presence of Gaussian noise. Comparing the results from 488 (61 × 8) samples as shown in Figure 1, 344 (43 × 8) samples in Figure 2, and 152 (19 × 8) samples in Figure 3, the errors and biases (and their spreads) increase as the number of samples is decreased as expected. In the case of 152 samples and at the lowest level of noise (*σ*_*g*_ = 0.0014, Lebedev, Knutsson, SH (L = 4 and 6), MAPL for shelled and non-shelled samplings with *N*_max_ = 6 lead to significant biases.

**Figure 2.**
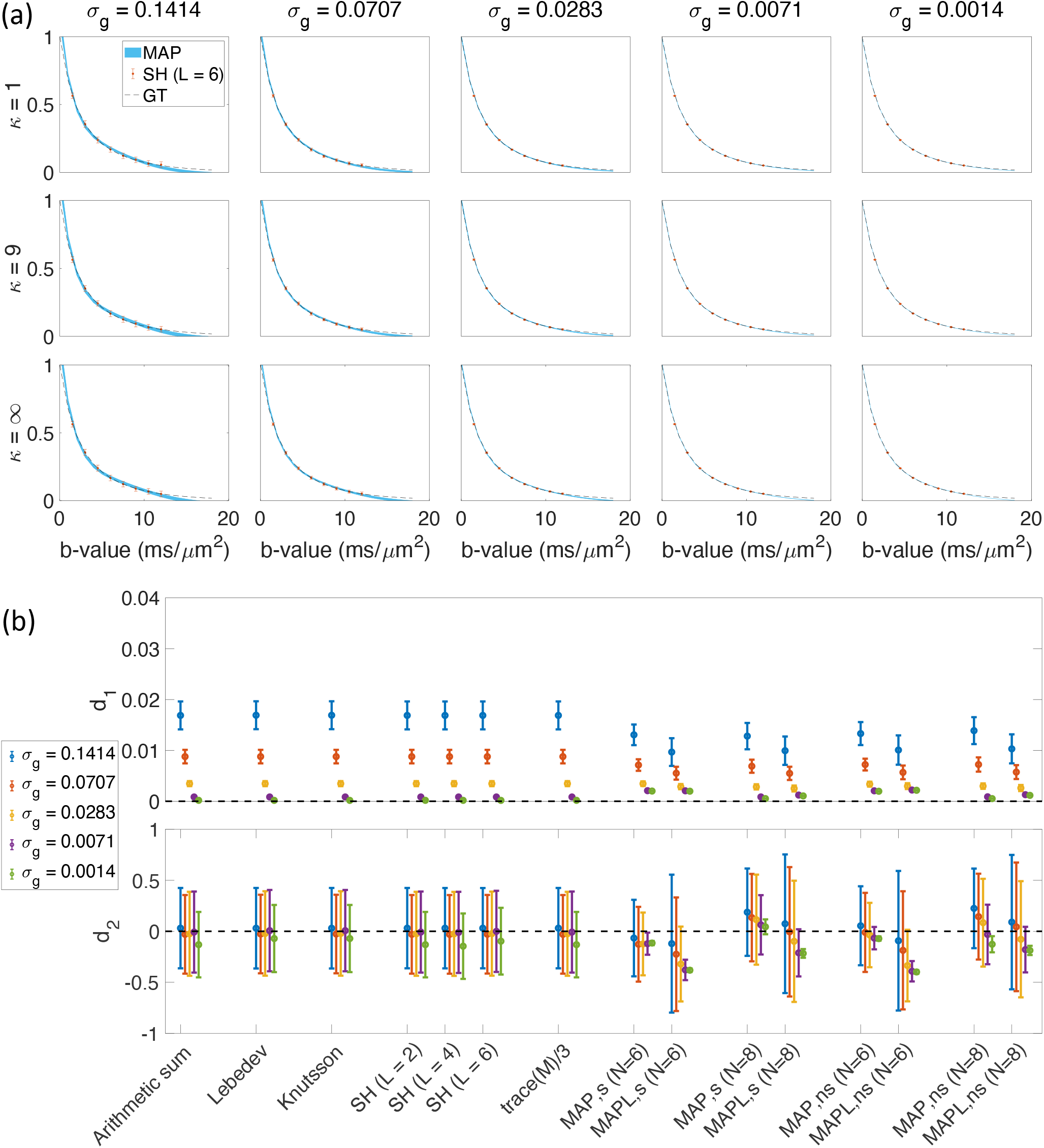
The results of 344 samples for both shelled (43 × 8)^26^ and non-shelled^21^ point sets in the presence of Gaussian noise. (a) the mean and std of the estimated signal versus b-value using MAP-MRI method with *N*_max_ = 6 for five different noise floors, *σ*_*g*_, and three different dispersion values, *κ*. The thickness of the blue band is twice the standard deviation of the signal estimates and its center is the mean. The dashed black line shows the ground truth and the red dots and bars show the results of the SH (L = 6), spherical harmonic representation. (b) the mean and std of the *d*_1_ and *d*_2_ measures for different methods. The ‘s’ and ‘ns’ correspond to the shelled and non-shelled point sets, respectively. Different colors (blue, red, yellow, purple, green) show the results in different noise levels (*σ*_*g*_ = 0.1414, 0.0707, 0.0283, 0.0071, 0.0014).

**Figure 3.**
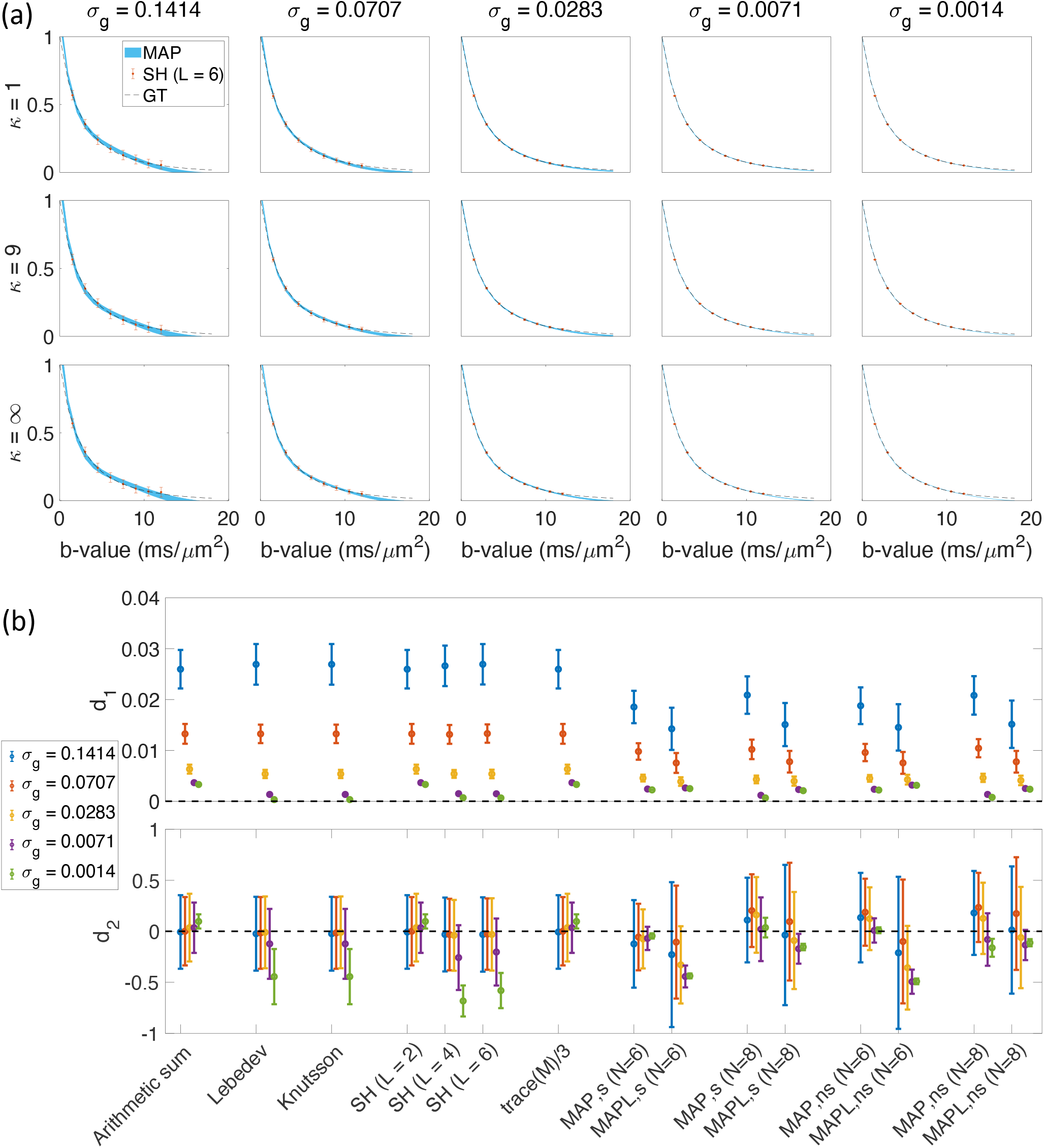
The results from 152 samples for both shelled (19 × 8)^26^ and non-shelled^21^ point sets in the presence of Gaussian noise. (a) the mean and std of the estimated signal versus b-value using MAP-MRI method with *N*_max_ = 6 for five different noise floors, *σ*_*g*_, and three different dispersion value, *κ*. The thickness of the blue band is twice the standard deviation of the signal estimates and its center is the mean. The dashed black line shows the ground truth and the red dots and bars show the results of the SH (L = 6), spherical harmonic representation. (b) the mean and std of the *d*_1_ and *d*_2_ measures for different methods. The ‘s’ and ‘ns’ corresponds to the shelled and non-shelled point sets, respectively. Different colors (blue, red, yellow, purple, green) show the results in different noise levels (*σ*_*g*_ = 0.1414, 0.0707, 0.0283, 0.0071, 0.0014).

Figure 4 shows the results obtained from the 344 (43 × 8) sample scenario in the presence of Rician signal. Clearly, when magnitude data is used and no bias correction scheme is employed, the noise will create a very significant bias (*d*_2_) in the signal that needs to be corrected^23^. Doing so would also help one obtain a reduction in the error, *d*_1_, in the estimation of powder average signal. Thus, we based our conclusions on the results obtained from simulated data assuming that the magnitude signal is transformed into Gaussian prior to the estimation of the orientational averages^24^.

**Figure 4.**
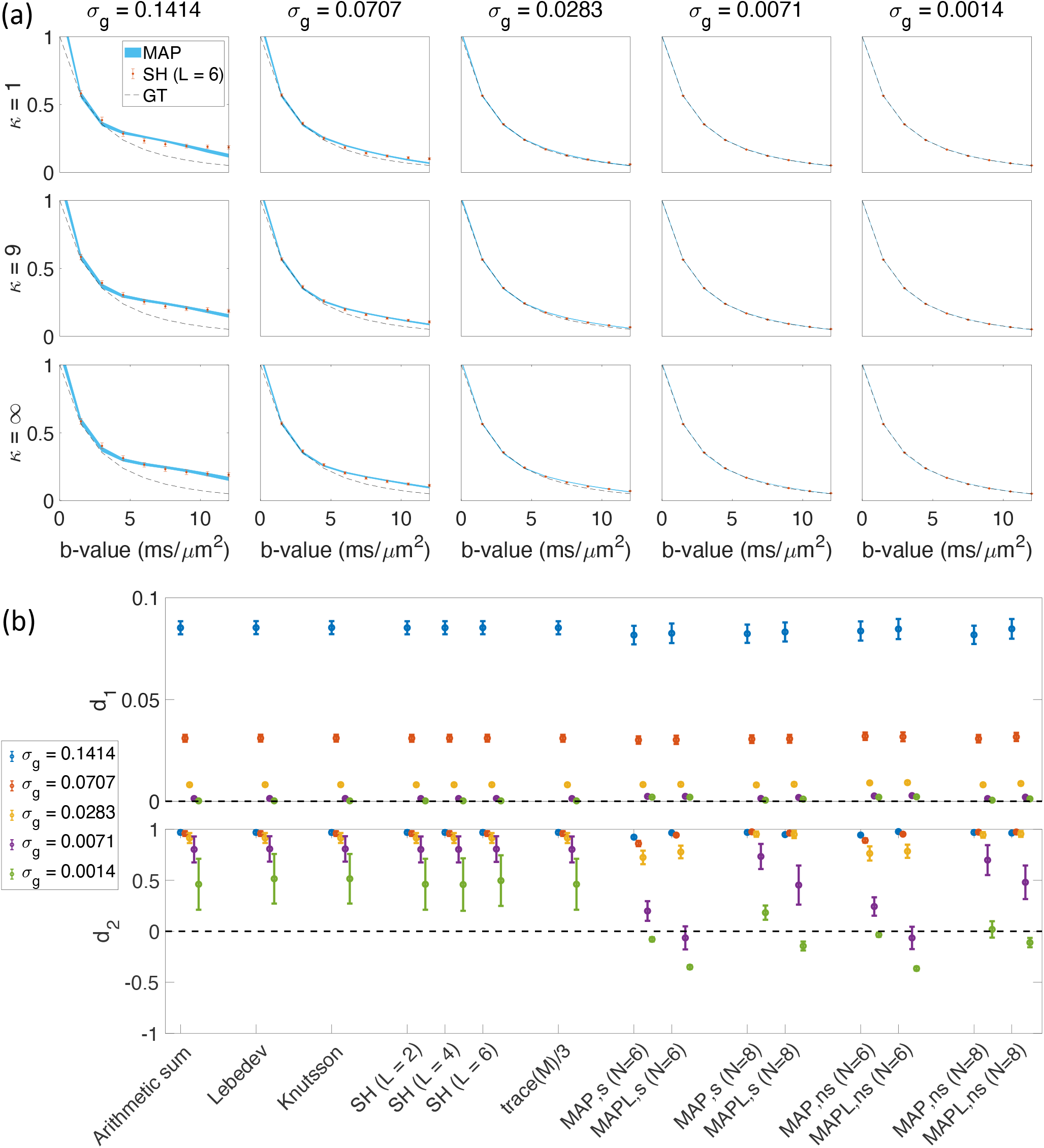
The results of 344 samples for both shelled (43 × 8)^26^ and non-shelled^21^ point sets for Rician signal. (a) the mean and std of the estimated signal versus b-value using MAP-MRI method with *N*_max_ = 6 for five different noise floors, *σ*_*g*_, and three different dispersion value, *κ*. The thickness of the blue band is twice the standard deviation of the signal estimates and its center is the mean. The dashed black line shows the ground truth and the red dots and bars show the results of the SH (L = 6), spherical harmonic representation. (b) the mean and std of the *d*_1_ and *d*_2_ measures for different methods. The ‘s’ and ‘ns’ correspond to the shelled and non-shelled point sets, respectively. Different colors (blue, red, yellow, purple, green) show the results in different noise levels (*σ*_*g*_ = 0.1414, 0.0707, 0.0283, 0.0071, 0.0014).

### Effect of dispersion

In Figure 5, we assessed the effect of dispersion on the estimates. We used the set of 344 (43 × 8) points and estimated *d*_1_ and *d*_2_ measures for Knutsson, MAP and MAPL (*N* = 6 and 8). When *κ* is large, there is little dispersion while decreasing *κ* increases the dispersion. For different *κ* values, the error (*d*_1_) and bias (*d*_2_) are about the same with slight improvement in *d*_2_ as *κ* goes down.

**Figure 5.**
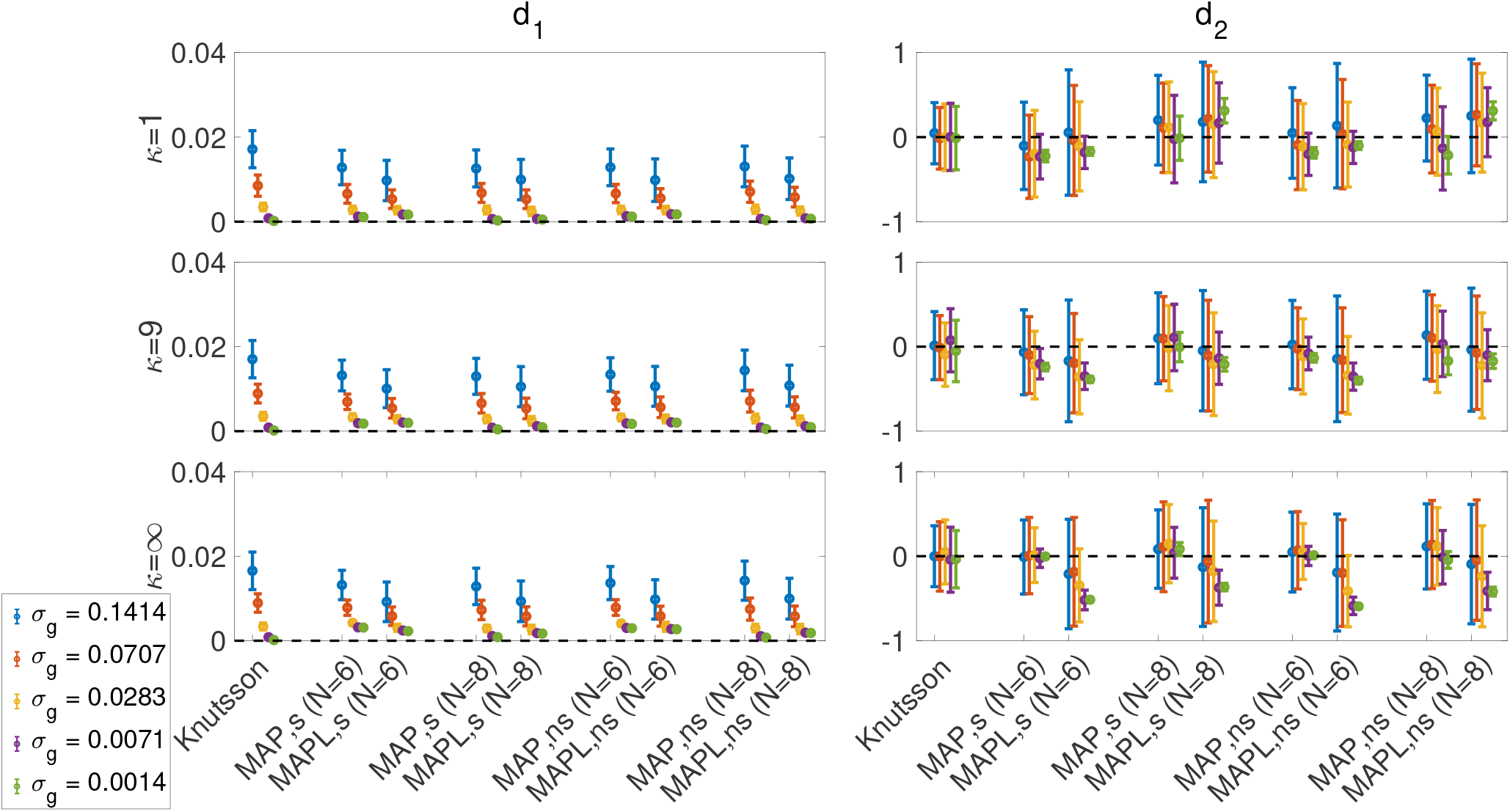
The estimated *d*_1_ and *d*_2_ for three different *κ* values, 344 (43 × 8) point sets in the presence of five different Gaussian noise levels. Note that when *κ* = ∞ there is no dispersion; decreasing *κ* increases the dispersion. For *κ* = 1, the amount of error, *d*_1_, and bias, *d*_2_ is smaller than *κ* = ∞ where there is no dispersion.

### Using MAP-MRI for interpolation

Figure 6 illustrates the results of MAP-based interpolation. We used both shelled (43 × 8) and non-shelled 344 point sets. To generate the ‘MAP, Knutsson, s8’ and ‘MAP, Knutsson, s43’, we utilized the shelled point sets. First, the signal was orientationally-averaged using the Knutsson method^25^ at *b* = 1.5, 3, 4.5, …, 12 *ms/μm*^2^, then the averaged signal was used to estimate the MAP-MRI coefficients in equation (19), i.e., we used the orientationally-averaged signal using Knutsson’s method as 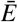 to estimate *κ*_(1+*N/*2)00_ in equation (19). Estimated coefficients (*κ*_(1+*N/*2)00_) are utilized to reconstruct the signal at *b* = 1.5, 3, 4.5, …, 12 *ms/μm*^2^ (‘MAP, Knutsson, s8’) and *b* = 1.5, 1.75, 2, …, 12 *ms/μm*^2^ (‘MAP, Knutsson, s43’). Note that ‘s8’ and ‘s43’ refer to the shelled point sets with 8, and 43 b-values, respectively, and number ‘43’ in ‘s43’ is independent from the number of gradient directions in 43 8 point set.

**Figure 6.**
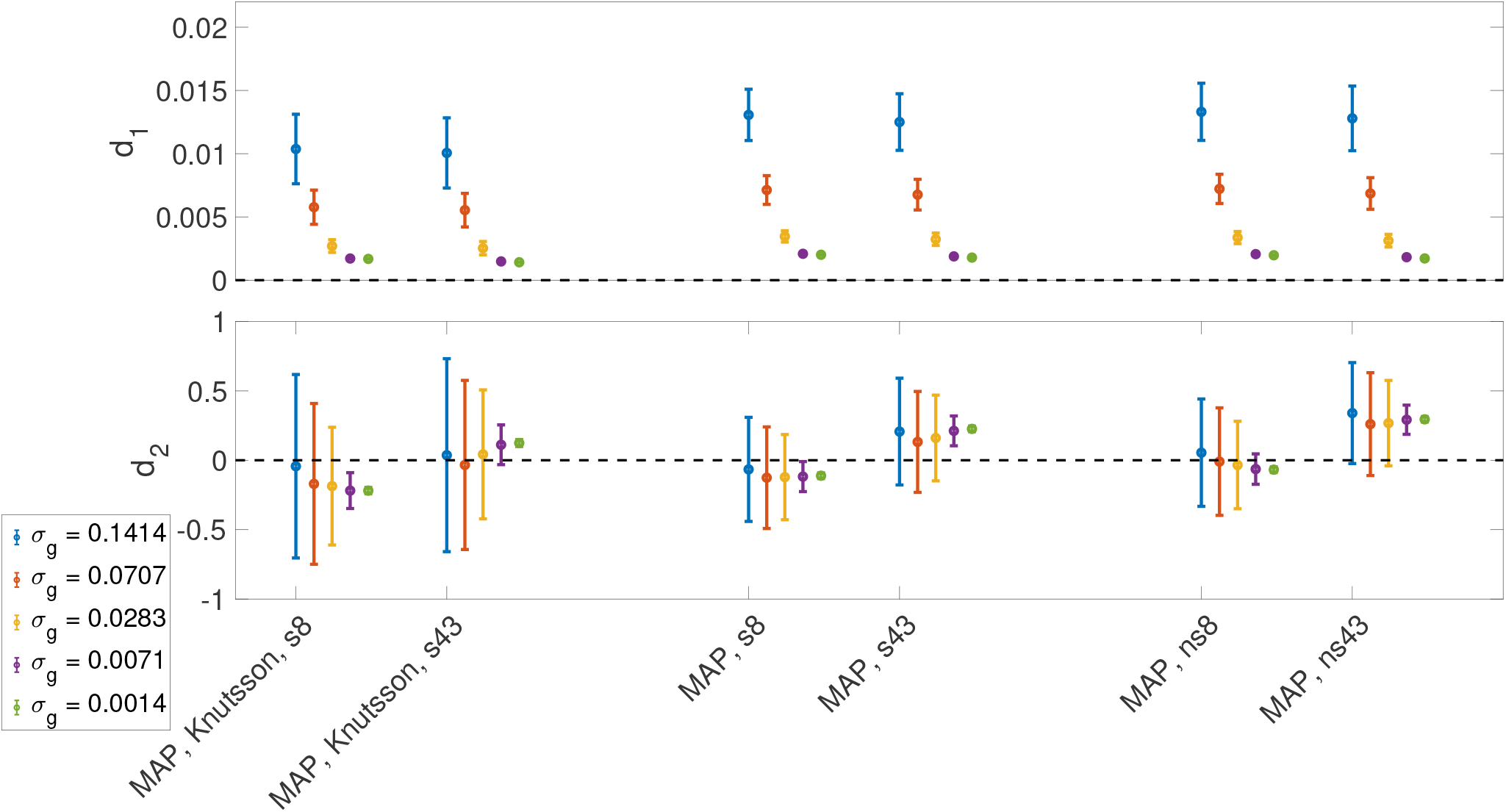
The results of MAP-based interpolation of the orientationally-averaged data from Knutsson method on 43 8 shelled Lebedev^26^ and non-shelled point sets^21^. The mean and std of the *d*_1_ and *d*_2_ measures for different scenarios. To generate the ‘MAP, Knutsson, s8’ and ‘MAP, Knutsson, s43’, we utilized the shelled point sets. First the signal is orientationally-averaged using Knutsson method^25^ at *b* = 1.5, 3, 4.5, …, 12 *ms/μm*^2^, then the averaged signal is used to estimate the MAP-MRI coefficients in equation (19), i.e., we use the orientationally-averaged signal using Knutsson method as 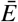 to estimate *κ*_(1+*N/*2)00_ in equation (19). Estimated coefficients (*κ*_(1+*N/*2)00_) are utilized to reconstruct the signal at *b* = 1.5, 3, 4.5, …, 12 *ms/μm*^2^ (‘MAP, Knutsson, s8’) and *b* = 1.5, 1.75, 2, …, 12 *ms/μm*^2^ (‘MAP, Knutsson, s43’). To generate ‘MAP, s8’ and ‘MAP, s43’ the shelled point set is used similar to the previous scenario but in this case we do not average the signal, all points are utilized to generate the MAP-MRI coefficients in equation (17) (*κ*_*jlm*_). Estimated coefficients (*κ*_*jlm*_) are utilized to reconstruct the signal at *b* = 1.5, 3, 4.5, …, 12 *ms/μm*^2^ (‘MAP, s8’) and *b* = 1.5, 1.75, 2, …, 12 *ms/μm*^2^ (‘MAP, s43’). Similar technique is used to generate the results of ‘MAP, ns8’ and ‘MAP, ns43’ for the non-shelled set of 344 directions. Note that when the MAP-MRI coefficients are used to estimate the powder average signal of *b* = 1.5, 3, 4.5, …, 12 *ms/μm*^2^ the results are the same as the ones reported in Figure 2. Using MAP-MRI for interpolation (*b* = 1.5, 1.75, 2, …, 12 *ms/μm*^2^) provides similar *d*_1_ and *d*_2_ compared to *b* = 1.5, 3, 4.5, …, 12 *ms/μm*^2^. Note that the *N*_max_ used in all the experiments in this figure is equal to 6.

To generate ‘MAP, s8’ and ‘MAP, s43’ the shelled point set was used similar to the previous scenario, but with the distinction that we do not average the signal; all points are utilized to generate the MAP-MRI coefficients in equation (17) (*κ*_*jlm*_). Estimated coefficients (*κ*_*jlm*_) are utilized to reconstruct the signal at *b* = 1.5, 3, 4.5, …, 12 *ms/μm*^2^ (‘MAP, s8’) and *b* = 1.5, 1.75, 2, …, 12 *ms/μm*^2^ (‘MAP, s43’). The same scheme is used to generate the results of ‘MAP, ns8’ and ‘MAP, ns43’ for the non-shelled set of 344 samples. The *N*_max_ used in all the experiments of this section (MAP-based interpolation) was taken to be 6.

The mean and std of the *d*_1_ and *d*_2_ measures for different methods (‘MAP, Knutsson, s8’, ‘MAP, Knutsson, s43’, ‘MAP, s8’, ‘MAP, s43’, ‘MAP, ns8’, and ‘MAP, ns43’) are illustrated in Figure 6. Note that when the MAP-MRI coefficients were used to estimate the powder-averaged signal of *b* = 1.5, 3, 4.5, …, 12 *ms/μm*^2^ the results were the same as those reported in Figure 2. Using MAP-MRI for interpolation (*b* = 1.5, 1.75, 2, …, 12 *ms/μm*^2^) provided similar *d*_1_ and *d*_2_ compared to *b* = 1.5, 3, 4.5, …, 12 *ms/μm*^2^.

## Discussion

Our analyses demonstrate that both the q-space sampling scheme and the numerical method for powder-averaging affect the estimated orientationally-averaged signal.

Our comparisons illustrated the relative performance of various methods subject to different noise levels and sampling schemes. Although our simulations are limited in terms of the complexity of the signal, we believe the simulated scenario is representative of the commonly encountered signal profiles. The analyses can be repeated in a manner similar to our approach for investigations targeting specific features (e.g. power laws) in the signal.

Our description of the methods for averaging the signal on a single shell illustrated that methods based on taking the ‘isotropic component’ of the signal yield weighted-averages of the original signal samples. The corresponding weights are given through equations (8)–(9) and (13). Among the methods that can be performed on a shell-by-shell basis, the Lebedev quadrature^26^ and Knutsson’s approach^27^ both show improved accuracy in the powder-averaged signal when compared to more simple arithmetic averaging approach, especially when there is a low number of point sets and the noise level is high.

The magnitude-valued MRI data is Rician distributed and our simulations show that without correcting for Rician noise, the powder-averaged signal is far from the ground truth especially when signal-to-noise ratio (SNR) is low. The SH and trace(M)/3 approaches lead to similar results as arithmetic averaging, with the difference that the SH approach outperforms arithmetic averaging when *L* = 4 or *L* = 6, especially for the scenario involving 19 directions.

We also investigated the bias introduced in the powder-average estimates, which can be relevant even in very high SNR scenarios. Various works in the literature, fitting multi-compartment models to derive microstructural parameters, have relied on information captured in the powder-averaged signal (e.g.^16, 28^). Employing an optimal q-space sampling scheme and proper direction-averaging method could significantly reduce the bias in parameters estimated from these models.

As gradient strengths are pushed higher and higher^29, 30^ maintaining gradient linearity becomes more and more challenging^31^, and truly multi-shell acquisitions become infeasible. In these situations, a representation of the diffusion MRI signal over the three-dimensional gradient sampling space becomes helpful. MAP-MRI is particularly well-suited to this situation as it provides a straightforward means of estimating the orientationally-averaged signal.

Here, we demonstrated that MAP-MRI-derived estimates can be used reliably, as they are more robust against noise compared to the shell-by-shell estimates of the orientational-average. The MAPL technique, which introduces regularization in lieu of constrained estimation, is preferred over the original MAP only when bias with b-values is not a concern. We note that the very recent formulation^32^ of the MAP method with hard constraints on the positivity of the estimated propagator (MAP+) could further improve the accuracy of the estimates.

## Methods

In this Section, we review the details of six different approaches to estimating the powder-averaged diffusion-weighted MRI signal.

### Arithmetic averaging

The simplest technique for powder-averaging is to distribute the samples as uniformly as possible over a sphere, and then compute the arithmetic mean of those measurements on that particular sphere^6^. In this scheme, the powder averaged signal for a given b-value 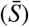 is estimated through

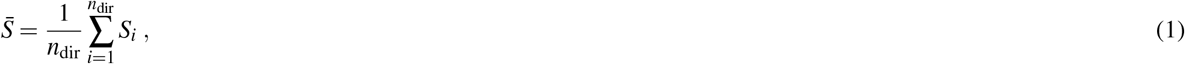

where *n*_dir_ is the number of gradient directions and *S*_*i*_ is the signal along the *i*th gradient direction. However, perfectly uniformly-distributed sets of directions are not readily attainable and therefore other methods have been proposed to overcome this problem.

### Weighted averaging

In most diffusion-weighted sampling schemes, weighted averaging can be used to account for the non-uniformity of the gradient directions. For each b-value, the powder averaged signal 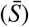 is

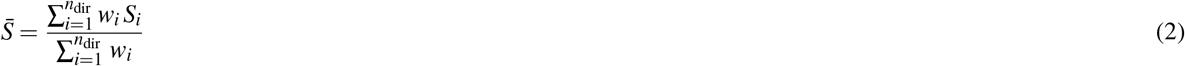

where *w*_*i*_ are the weights corresponding to the signal along the *i*th gradient direction. We considered four techniques that enable the determination of an optimal set of weights, *w*_*i*_, for the estimation of the orientationally-averaged signal.

#### Quadratures on the sphere (Lebedev)

Lebedev quadrature evaluates integrals over the unit sphere, i.e.,

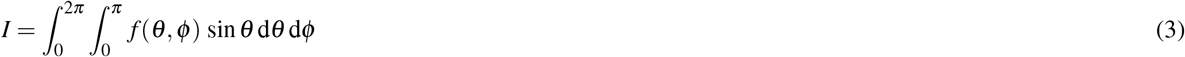

as an approximation

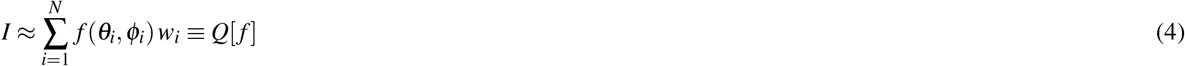

where the points (*θ*_*i*_, *φ*_*i*_) and weights *w*_*i*_ are estimated for different *N* using the algorithm in^26^. Q[f] is the quadrature of the exact integral. Similar to the one-dimensional Gauss quadratures, the nodes, (*θ*_*i*_, *φ*_*i*_), and the weights can be determined simultaneously. A constraint is imposed to integrate all spherical harmonics up to degree *p*. This leads to a system of nonlinear equations, which can be solved to provide the optimal nodes and weights. This idea stems from the work by Sobolev^33^.

Lebedev built a set of quadratures and the set of non-linear equations are solved for degrees up to *p* = 131^34–38^, yielding one of the most commonly-used quadratures for integration over the sphere.

In this study, we employ this technique by setting *f* (*θ, φ*) to be the signal profile at a particular *b*-value. As this technique requires the value of the function to be evaluated along specific directions, the set of gradient directions has to be chosen accordingly, i.e., the technique cannot be readily employed with commonly-available diffusion MRI sampling protocols.

#### Representation of the single-shell signal in a series of spherical harmonics

Any square-integrable function, 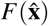, defined over the unit sphere, can be represented in a spherical harmonic basis through the expansion

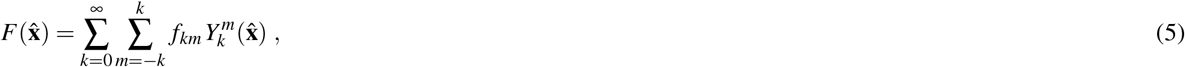

where *k* and *m* indicate the order of the spherical harmonics function denoted by 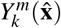. The coefficients are given by

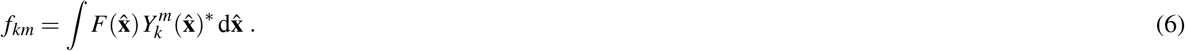

If 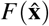) is rotationally-invariant, i.e., independent of 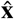, all coefficients except *f* _00_ vanish.

Spherical harmonics can be used to represent the diffusion signal profile at a fixed *b*-value^39^ by terminating the series at a finite value *k*_max_ = *L*. In matrix form, such a representation can be written as

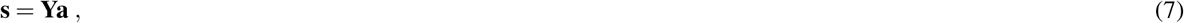

where 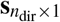 is the vector of diffusion-weighted signals, 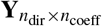 is the matrix of spherical harmonics for *n*_coeff_ = (*L* + 1)(*L* + 2)*/*2 number of coefficients, and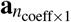 is the vector of coefficients, which can be estimated using

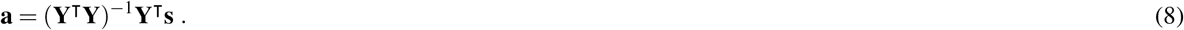

As mentioned earlier, an estimate of the powder-averaged signal is obtained by taking the signal’s “isotropic component” in its irreducible representation^3^. When adopted to SDE acquisitions on a single shell, the orientationally-averaged signal is given simply by the coefficient corresponding to the *k* = *m* = 0 term of the series, or more explicitly,

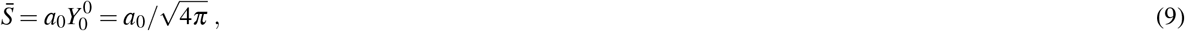

where we employ the convention that 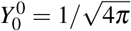^40^.

Note that equations (8) and (9) suggest that the powder-average estimate using this scheme is also a weighted average of the signal values, albeit without any constraint on the selection of gradient directions.

#### Representation of the single-shell signal using Cartesian tensors

The diffusion-weighted signal profile at a fixed b-value can also be represented by the following equation:

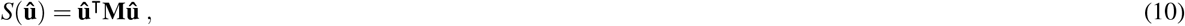

where 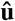 is the unit vector denoting the gradient direction and **M** is a 3 × 3 symmetric positive definite matrix. In this representation, the orientationally-averaged signal is given by

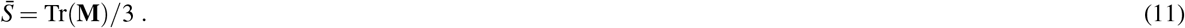

Considering that **M** is symmetric, we can cast the problem in the matrix form

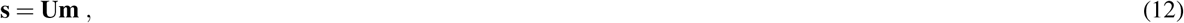

where 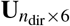 is the matrix whose each row is given in terms of the corresponding gradient direction 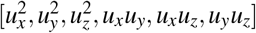, and **m**= [*M*_*xx*_, *M*_*yy*_, *M*_*zz*_, *M*_*xy*_, *M*_*xz*_, *M*_*yz*_].

The powder-averaged signal (11) can thus be estimated via

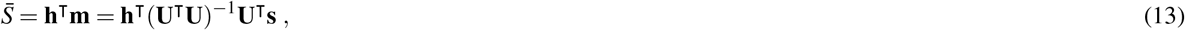

where **h**= [1*/*3, 1*/*3, 1*/*3, 0, 0, 0]^r^.

As discussed in the context of representing the apparent diffusivity profiles^41, 42^, from a conceptual point-of-view, the representation (10) is equivalent to the representation of the signal in terms of a series of spherical harmonics terminated at *k* = 2. However, we include it here as a separate scheme to assess potential numerical differences between the methods. We also note that representations in terms of higher order Cartesian tensors equivalent to the series of spherical harmonics terminated at higher orders can be formulated^41^, but excluded here for brevity.

#### Knutsson

Following from spherical harmonic representation of the signal, the rotational variance of a set of unit vectors 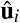, with *i* = 1 *… n*_dir_, can be analyzed using spherical harmonics in the following way^25, 27^. Consider a weighted sampling function of the form

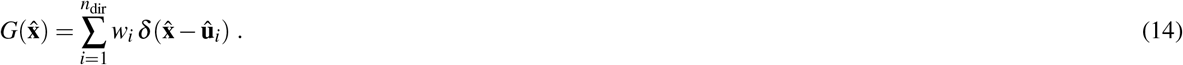

The coefficients of this function in the spherical harmonic basis is given by

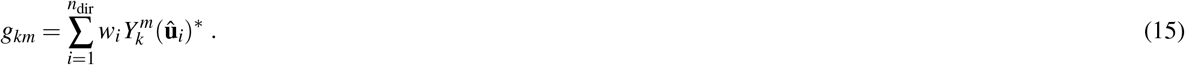

For rotationally-invariant sampling, these coefficients would obey *g*_*km*_ ∞ *δ*_*k*0_*δ*_*m*0_. Let us denote by **g**_0_ the vector of coefficients having non-zero value only when *k* = *m* = 0. An optimal vector of weights, **w**_0_, can be obtained using the expression^25^

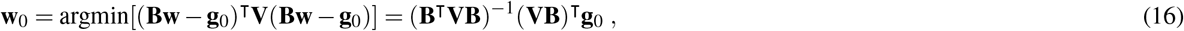

where **B** is the matrix of the spherical harmonics basis sampled at the orientation 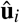, and **V** is a diagonal matrix containing the weights for each spherical harmonic^25^. **V**, which corresponds to 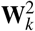 in equation (4) in^25^, is used to set the importance of obtaining the specified response for different spherical harmonic basis functions which relate to the expected signal content of different spherical harmonics in the measurement. This depends on the tissue as well as on the b-value. We found that taking **V** to be a diagonal matrix with diagonal elements given by (1 + *k*^2^*/*36)^−1^ provides adequate distribution of weights to respective terms of the series for typical signal decay profiles within the brain parenchyma. The size of the matrices depended on the number of directions. In the implementation, the maximum order *k* was taken to be 18, 14, and 10 for 61, 43, and 19 directions, respectively.

We note that using this procedure, arithmetic averaging could be justified only for those sets of unit vectors that lead to equal weights, *w*_*i*_. Such sets of unit vectors are not attainable for all but a few very special cases of *n*_dir_. Equation (16) and the result of spherical harmonic representation are the same (with **B**= **Y** r) if the weighting function **V** in equation (16) is unity for degree 1 to *L* and zero for higher *L*.

### MAP-MRI

Above, we described several techniques with which the orientationally-averaged signal can be estimated from single-shell data wherein the data points are collected by repeatedly applying gradients along different directions while keeping the b-value fixed. In most applications involving the powder-averaged signal 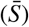, one is interested in characterizing the dependence of the signal on the b-value. This can be accomplished by acquiring data on multiple shells and repeatedly applying one of the above methods on each shell. Alternatively, one can employ a representation of the signal on the entire three-dimensional space sampled by the gradient vector and compute its “isotropic component” similar to what was done above for the spherical harmonics representation.

MAP-MRI^22^ is a powerful representation, which was shown to not only reproduce the diffusion-weighted signal over the three-dimensional space (see the result of ISBI challenge 2020 as an example,^43, 44^ but also provide accurate estimates of scalar measures from datasets including heavily diffusion-weighted acquisitions^45^. One scalar index derived from such sampling is the propagator anisotropy (PA) whose formulation involves the estimation of the isotropic component of the propagator^22^, which can be adopted for estimating the orientationally-averaged signal as indicated before. This is accomplished by using the formulation of MAP-MRI in spherical coordinates^22, 46^. In Cartesian coordinates, the formulation of MAP-MRI follows, in a straightforward manner, from its one-dimensional version^47^ that features Hermite polynomials and allows for anisotropic scaling of the basis functions. When expressed in spherical coordinates, the following equation is used for this purpose (equation (58) in^22^):

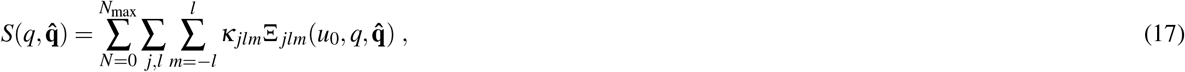

where *j* ≥ 1, *l* ≥ 0 and 2 *j* + *l* = *N* + 2, *u*_0_ is a scalar related to the width of the basis functions, and *q* and 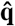 are the magnitude and direction of the wavevector, which is proportional to the gradient vector. The basis function 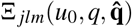 is given by

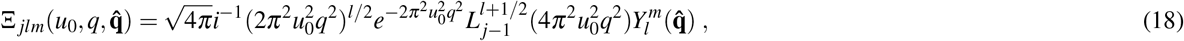

where 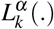 is the associated Laguerre polynomial and 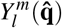 is the spherical harmonic.

The isotropic part of the diffusion-weighted signal is the powder-averaged signal, which is obtained by setting *l* = *m* = 0 and *j* = 1 + *N/*2:

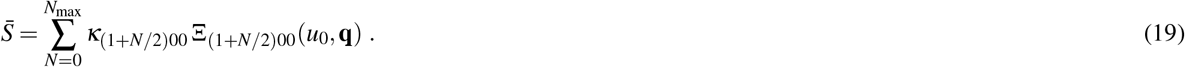

In this study, we employed two versions of the MAP-MRI technique: (i) its original formulation by Özarslan et al., in which the positivity of the propagator is enforced over a large domain in displacement space^22^; and (ii) a later formulation called MAPL introduced by Fick et al.^48^ in which a Laplacian regularization is employed instead of the constraints, similar to what was done in the corresponding one-dimensional problem^49^.

Table 1 summarizes the different powder-averaging techniques used in this study.

**Table 1.** The summary of different powder-averaging techniques used in this study.

|  | Estimation method |
| --- | --- |
| 1- Shell-by-shell estimation | 1- Arithmetic averaging<sup>6</sup><br>2- Quadratures on the sphere (Lebedev)<sup>26</sup><br>3- Representation of signal in spherical harmonic<sup>3</sup><br>4- Representation of signal in Cartesian tensors<br>5- Knutsson<sup>25</sup> |
| 2- Estimation from the entire 3D data | 1- MAP-MRI<sup>22</sup><br>2- MAP with Laplacian regularization<sup>48</sup> |

### Simulations

The noise-free diffusion-weighted signal at a b-value *b* when the gradients are applied along the unit vector 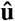 was generated using the following equation:

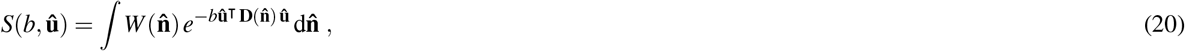

where 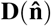 is an axisymmetric, prolate tensor oriented along 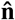 with eigenvalues *D*^||^ = 1 *μm*^2^*/ms*, and *D*^┴^ = 0.14 *μm*^2^*/ms*. The orientation distribution function, 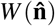 is taken to be a Watson distribution function given by

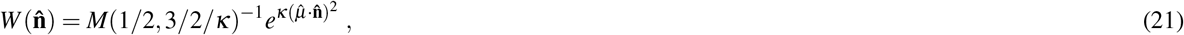

where *M* is the confluent hypergeometric function, 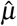 is the mean direction, taken to be (0.4, 0.6, 0.693)^r^ and *κ* is the concentration parameter. When *κ* is small, for example, *κ* = 1 we have high orientation dispersion (i.e, we have a fat orientation distribution function (ODF)) and when *κ* is large, for example, *κ* = 64, the orientation dispersion is small and the ODF is sharp. In our simulation, we used *κ* = 1, 9, and ∞ (a signal without dispersion or Dirac delta ODF). Irrespective of the orientation distribution function, the ground truth orientationally-averaged signal is given by^12, 50^

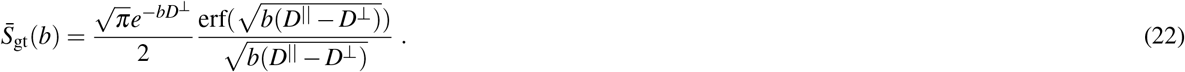

The multi-shell data were computed at the b-values *b* = 0, 1.5, 3, 4.5, 6, 7.5, 9, 10.5, and 12 *ms/μm*^2^ and with Δ = 43.1 and *δ* = 10.6 *ms*. For the above parameters and with *b* = *q*^2^(Δ − *δ /*3), we expect the following signal values:

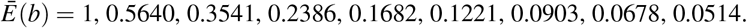

The noisy diffusion-weighted signal values are synthesised according to the following for Gaussian and Rician distributed signals, respectively:

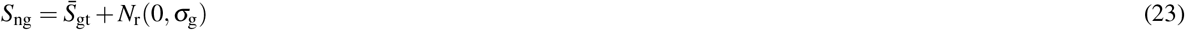

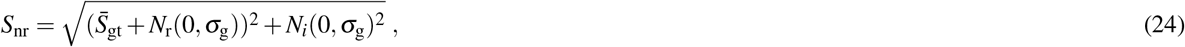

where *N*_*r*_(0*, σ*_g_) and *N*_*i*_(0*, σ*_g_) are the normal distributed noise in, respectively, the real and imaginary images with a standard deviation of *σ*_g_. Here, we simulate the noisy signal with *σ*_g_ = 0.1414, 0.0707, 0.0283, 0.0071, and 0.0014.

### Evaluation criteria

We utilized two measures *d*_1_ and *d*_2_ to quantify the fidelity of the orientationally-averaged signal. *d* _1_ shows the absolute error between the estimated signal and the ground truth. For the correlation between the bias, *∊*_*j*_, and the b-values, the Pearson’s correlation coefficient (*d*_2_) was used.

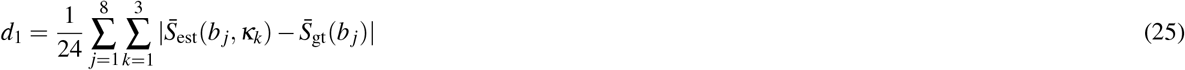

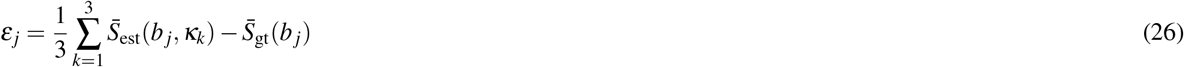

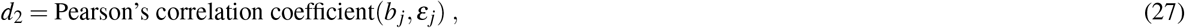

where 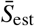 and 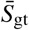 are, respectively, the estimated powder average signal and the ground truth values.

### Point sets

#### Shelled point sets

The gradient directions were generated using Knutsson’s method to produce 61 gradient samples^21^ and Lebedev’s method^26^ to produce sets with 43 and 19 directions. The signal values were estimated at 8 b-values along each direction. For each sampling scheme, the same number of *S*(0) signal values were considered (i.e. 61 *S*(0) signal values for Knutsson’s method^21^ and 43 and 19 *S*(0) signal values for Lebedev’s method^26^).

#### Non-shelled point sets

The three-dimensional (non-shelled) gradient vector sets were generated using Knutsson’s method for 488, 344, and 152 gradient directions^21^ matching the total number of samples (61 × 8, 43 × 8, and 19 × 8) in the shelled point sets. For 488, 344, and 152 sampling schemes 61, 43, and 19 *S*(0) signal values were considered.

Table 2 shows a summary of these point sets.

**Table 2.** The summary of different vector sets and different experiments conducted in this study.

| Point sets | Sampling scheme | Reference |
| --- | --- | --- |
| shelled | 488 (61×8) samples (Knutsson) | 21 |
|  | 344 (43×8) samples (Lebedev) | 26 |
|  | 152 (19×8) samples (Lebedev) | 26 |
| non-shelled | 488 samples (Knutsson) | 21 |
|  | 344 samples (Knutsson) | 21 |
|  | 152 samples (Knutsson) | 21 |
| Experiments | Sampling scheme |  |
| Effect of noise and number of samples | 488 (61×8) samples<sup>21</sup> with Gaussian noise | Figure 1 |
|  | 344 (43×8)<sup>21,26</sup> samples with Gaussian noise | Figure 2 |
|  | 152 (19×8)<sup>21,26</sup> samples with Gaussian noise | Figure 3 |
|  | 344 (43×8)<sup>21,26</sup> samples with Rician noise | Figure 4 |
| Effect of dispersion | 344 (43×8) samples<sup>21,26</sup> with Gaussian noise | Figure 5 |
| Using MAP-MRI for interpolation | 344 (43×8) samples<sup>21,26</sup> with Gaussian noise | Figure 6 |

### Experiments

We conducted three different experiments to investigate the effect of encoding scheme and powder-averaging technique on the estimated orientationally-averaged signal.

#### Effect of noise and number of directions

For designing an experiment, one of the important factors is minimizing the total acquisition time (for in vivo applications). However, at the same time, the set of measurements should provide sufficient information for robust model fitting. The acquisition should therefore be optimized to provide maximum information per unit time. In this work, we compared three different sets of sampling vectors, 61 × 8, 43 × 8, and 19 × 8, (shelled and non-shelled) with different noise levels to investigate the effect of encoding scheme and noise on the estimated powder-averaged signal.

#### Effect of dispersion

The powder-averaged signal should provide rotationally-invariant tissue measures. In addition to the sampling scheme mentioned in the previous section the amount of actual orientational dispersion in the underlying structure will also affect the accuracy of the orientionally-averaged signal estimates.

#### Using MAP-MRI for interpolating the orientationally-averaged signal

In recently published works that employ the orientationally-averaged signal^10,13,16,28^, the acquisition is performed using shell-based sampling schemes, and usually restricted by acquisition time. As a result, the number of shells is not much higher than the number of parameters in the model, which could result in inaccurate parameter estimates. The ability of MAP-MRI to estimate the signal values across the q-space can be used to address this challenge. To illustrate this, a set of coefficients is estimated from the measurements that can be used to provide the signal values for intermediate b-values that are not sampled. This can be especially useful when the orientationally-averaged signal is applied in multi-compartment models.

Table 2 summarizes different experiments conducted in this study.

## Acknowledgements

MA and DKJ are supported by a Wellcome Trust Investigator Award (096646/Z/11/Z) and DKJ by a Wellcome Trust Strategic Award (104943/Z/14/Z). This project was financially supported by the Swedish Foundation for Strategic Research (AM13-0090 and Multimodal Guidance in Neurosurgery), the Swedish Research Council 2015-05356 and 2016-04482, Linköping University Center for Industrial Information Technology (CENIIT), LiU Cancer, VINNOVA/ITEA3 17021 IMPACT, and Analytic Imaging Diagnostic Arena (AIDA).

## Author contributions statement

EÖ and MA conceptualized the study. MA performed research with inputs from EÖ and HK. DKJ provided guidelines and funding. All authors collaborated on writing and editing the manuscript.

